# LED-based interference-reflection microscopy combined with optical tweezers for quantitative three-dimensional single microtubule imaging

**DOI:** 10.1101/277632

**Authors:** Steve Simmert, Mohammad Kazem Abdosamadi, Gero Hermsdorf, Erik Schäffer

**Affiliations:** Cellular Nanoscience, Center for Plant Molecular Biology, University of Tübingen, Auf der Morgenstelle 32, 72076 Tübingen, Germany

## Abstract

Optical tweezers combined with various microscopy techniques are a versatile tool for single-molecule force spectroscopy. However, some combinations may compromise measurements. Here, we combined optical tweezers with total-internal-reflection-fluorescence (TIRF) and interference-reflection microscopy (IRM). Using a light-emitting diode (LED) for IRM illumination, we show that single microtubules can be resolved with high contrast. Furthermore, we converted the IRM interference pattern of an upward bent microtubule to its three-dimensional (3D) profile calibrated against the optical tweezers and evanescent TIRF field. In general, LED-based IRM is a powerful method for high-resolution 3D microscopy.

**OCIS codes:** (180.3170) Interference microscopy; (120.4570) Optical design of instruments (350.4855); Optical tweezers or optical manipulation.

## 1. Introduction

To apply force on a wide range of samples and measure the response down to single biological molecules, optical tweezers are widely used in physics, material science, and biology [1]. In the optical trap, the momentum of light is used to apply forces onto trapped cells or particles. For example, microspheres are often used as sensor probes to measure the mechanochemical interaction of individual molecular machines with their tracks or substrate. In case of cytoskeletal motor proteins, either actin filaments or microtubules are immobilized on surfaces [1]. During an experiment, such filaments need to be imaged for extended periods of time. To this end, optical tweezers are often combined with various microscopy contrast methods ranging from bright-field, via differential interference contrast (DIC) [2, 3], or less common, phase-contrast [4] and dark field [5] to fluorescence including among others epi-, TIRF, confocal, or STED configurations [6–10]. Because fluorescence microscopy requires labeling of proteins, is phototoxic, and has photobleaching, and brightfield microscopy has low contrast or involves considerable post-processing of the images [11], the technique of choice for visualizing microtubules is often video-enhanced DIC [2, 12]. However, due to the shear axis of the Nomarski prisms, simple DIC implementations are direction-dependent [13] and limit the polarization and power of optical tweezers. Only linearly polarized trapping lasers with the polarization direction aligned with the Nomarski prism’s shear axis pass the prims without changing the polarization. Linearly polarized trapping light induces an asymmetry in the force field of the optical trap [14], which may lead to artifacts for 2D and 3D experiments [15, 16], and makes experiments that utilize the transfer of angular momentum of the trapping light to trapped particles difficult [17]. In addition, for near infrared trapping lasers, about 10 % of power is lost per prism. Despite these limitations, DIC is often the technique of choice for microtubule-based, optical-tweezers assays.

To eliminate these restrictions, we combined our optical tweezers with IRM. In addition, we integrated TIRF into the system to allow for single-molecule fluorescence microscopy. IRM provides high, three-dimensional (3D) contrast, can be realized with a simple, cost-efficient optical design, and, in comparison to differential interference contrast (DIC), is orientation independent and does not compromise the polarization state or the power of optical tweezers. Furthermore, since IRM works in reflection, the half-space above the objective is free for other manipulation techniques. IRM was first used by Curtis *et al.* [18] to measure cell-substrate distances during cell adhesion. These distances were inferred from interference fringes that occur because the back-scattered (or reflected) light of the specimen interferes with the reflected “reference” light from the glass–water interface converting a phase into an amplitude contrast. This contrast was improved significantly by reducing the amount of unwanted reflected light from other optical elements by crossed polarizers and a so-called “Antiflex” objective (reflection interference contrast microscopy (RICM) [19]). Furthermore, a quantitative analysis of separation distances and optical thicknesses is possible [20] providing 3D information about the sample with exquisite axial resolution allowing, for example, ångström precision tracking [21]. Recently, the detection limits of the technique have been pushed further, utilizing the technical improvements of digital cameras, processing power, and lasers for illumination (interferometric scattering microscopy (iSCAT) [22, 23]) allowing for the detection of single unlabeled molecules [24, 25]. Single microtubules were imaged by IRM in a confocal configuration [26], more recently with high spatiotemporal resolution using iSCAT [27], or a solid-state-white-light illumination in an inverted epi-fluorescence microscope [28].

In our combination with optical tweezers, we used a high-power LED as a light source and show that single microtubules can be visualized with a signal-to-noise (SNR) ratio comparable to DIC. We measured and optimized the microtubule contrast in terms of photon-flux, frame-averaging, and illumination numerical aperture. Furthermore, using the optical tweezers and the height-dependent interference patterns of an upward bent microtubule, we measured its 3D profile. We used this profile to calibrate the evanescent field of the TIRF microscope. The short coherence length of the LED compared to lasers has the advantage that no speckles and etalon fringes arise, which simplifies image analysis procedures and improves contrast. In general, LED-based IRM is a simple, but powerful microscopy contrast method compatible with many other techniques allowing for high-resolution 3D microscopy. Our combination can be applied to a wide range of biological systems and allows versatile imaging and force spectroscopy with molecular resolution.

## 2. Materials and methods

### 2.1. Experimental setup

The setup combines optical tweezers with TIRF microscopy and IRM (Fig.1) described in more detail below. The assembly is custom-built and placed in a vibration-isolated walk-in chamber inside an air-conditioned room. The temperature of the chamber itself is not actively controlled, but isolates the device from outside noise and heat sources. All components that generate heat are constantly on such that the room equilibrates to a constant temperature. Chambers are only entered to exchange samples. During an experiment, all adjustments are done via motorized components. All components are placed on an 1.2 m × 1 m × 0.2 m optical table (Standa, Lithuania), which is mechanically stabilized by an active vibration isolation system (VarioBasic-60, Accurion, Germany). Overall, the system had a tracking precision (Allan deviation) of a fixed microsphere of less than 5 nm and 10 nm for the lateral and axial direction, respectively, measured over a duration of 1000 s. Over a minute, the tracking precision was better than 1 nm in all directions. Thus, thermal drift was not a problem for the current study. In the following, we describe the optical tweezers, the IRM, and TIRF implementation.

**Fig. 1:**
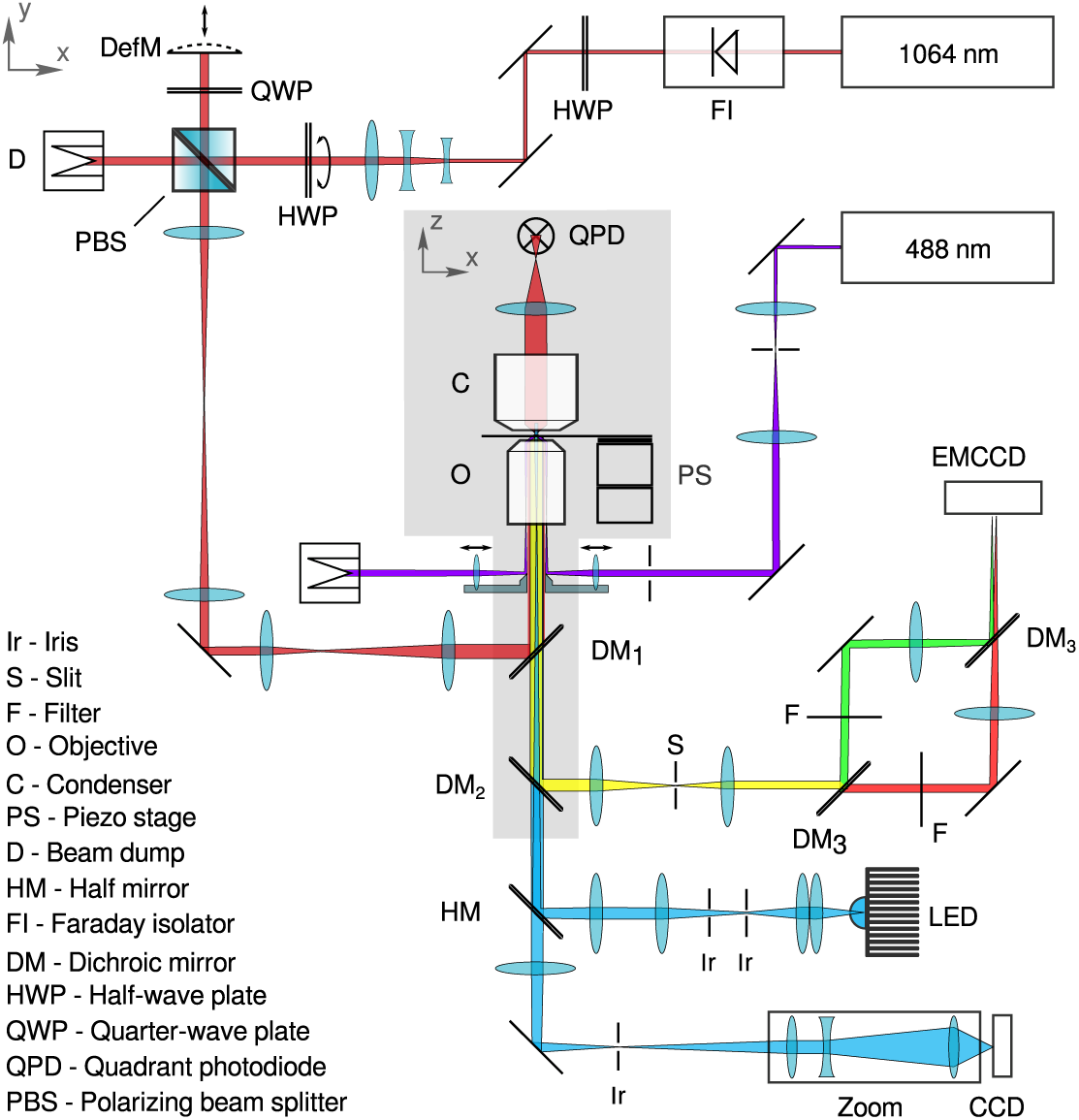
Schematic of the combined IRM–TIRF-microscope–optical-tweezers setup.

#### 2.1.1. Optical tweezers setup

The optical tweezers setup uses a diode-pumped solid state YVO_4_-laser that continuously emits 3 W at 1064 nm (Smart Laser Systems, Germany). The laser passes a Faraday isolator to avoid back-reflections and etalon fringes. The half-wave plate (HWP) matches the laser polarization with the following mirror axes to preserve the linear polarization of the trapping laser. The 3-lens Galilean telescope increases the beam diameter to 5 mm. Adjustment of the laser intensity is realized by a combination of a HWP and a polarizing beam splitter (PBS) cube. Unwanted laser light passes the PBS and reaches a beam dump, whereas the reflected light passes a quarter-wave plate (QWP) and is reflected by a deformable mirror (MMDM10-1-focus, OKO Technologies, The Netherlands). This mirror enables fast axial steering of the trap in the sample relative to the imaging plane of the microscope [29]. After the reflection, the laser passes the QWP a second time resulting in a 90^°^ rotation of the linear polarization direction. Thus, the laser is transmitted by the PBS and passes two Keplerian telescopes. A dichroic mirror (F33-725, AHF Analysentechnik, Germany) reflects the laser beam to the objective lens (CFI Apo TIRF 60x oil, NA 1.49, Nikon Instruments, Japan). The two telescopes relay the optical plane of the deformable mirror into the back-focal plane (BFP) of the objective lens and magnify the laser beam-diameter to 8.8 mm corresponding to an overfilling ratio of α = 1.0 [30]. The objective tightly focuses the laser and forms a trap in the sample chamber. The sample is mounted on a long-range open-loop (MX-35, Mechonics, Germany) and a short-range, high-precision, closed-loop piezo-driven stage (PI Hera 620 XYZ, Physik Instrumente, Germany). The transmitted laser light is collected by a condenser lens (D-CUO Achr.-Apl. NA 1.4, Nikon Instruments, Japan). The image of the BFP of the condenser is de-magnified and projected onto a quadrant photo diode (QP154-Q-HVSD, First Sensor AG, Germany) by a relay lens. To ensure thermal stability in the sample, both the objective and condenser lens are kept at 29.000 ^°^C by a temperature feedback control [3].

#### 2.1.2. Interference reflection microscope

The LED-based interference reflection microscope consists of a few simple optical components (Fig. 2). For illumination a blue LED (Royal-Blue LUXEON Rebel LED, Lumileds, Germany; λ ≈ 450 ± 20 nm with 525 mW at 700 mA) was used. To provide thermal stability—especially at high currents, the LED was mounted on a large aluminum heat sink (Fisher Elektronik, Germany). The driving current is provided by a DC power supply (Agilent E3648A, Keysight, Böblingen, Germany). An image of the LED is magnified by two telescopes and projected into the BFP of the objective lens to Köhler-illuminate the sample. An aperture iris is placed at the first image plane of the LED, which is conjugate to the BFP. This iris truncates the LED image and, thus, avoids total internal reflection. With the aperture iris, the illumination numerical aperture (INA) can also be adjusted. A field iris limits the illumination area in the sample-plane, i.e. the front-focal plane of the objective. The detection and illumination light path are separated by a 50/50 beam splitter plate (BS) (F21-000, AHF Analysentechnik, Germany). Half the reflected light from the sample passes the beam splitter. A 200 mm tube lens and a zoom (S5LPJ7073, Sill Optics, Germany) magnify the sample image about 165× and project it onto a CCD camera (LU135-M, Lumenera, Canada). Note that the use of an Antiflex objective and linearly polarized light did not improve the image contrast significantly.

**Fig. 2:**
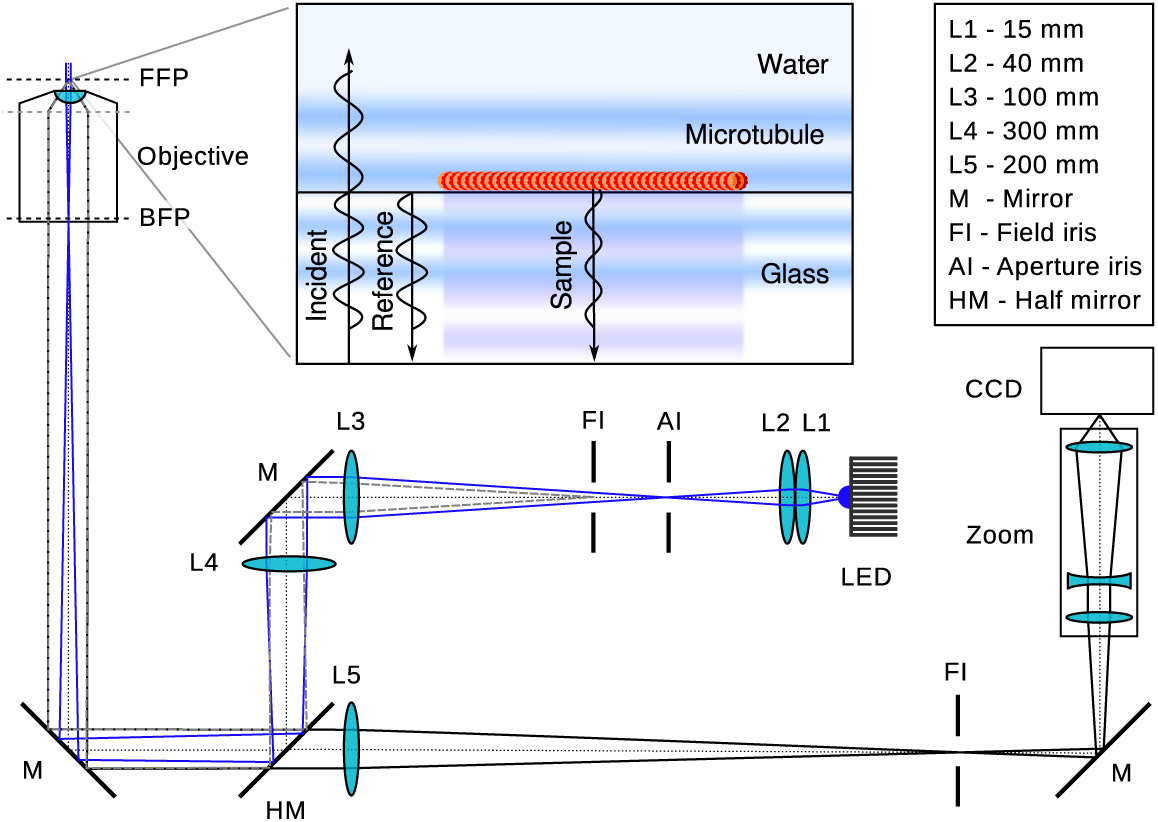
**Schematic of the IRM design and principle.** The optical path is drawn to scale and shows the marginal rays of the LED (blue lines), the sample (black lines), and the field iris (dashed gray lines). The inset shows the principle of the IRM contrast formation. The incident light of the Köhler-illuminated sample, here a microtubule, is reflected from the glass-water interface (reference) and the specimen (sample), which interfere to form the final signal. Note that the light fields are illustrated at normal incidence, which would correspond to an illumination numerical aperture (INA) close to zero. A higher INA leads to tilted wavefronts omitted for clarity.

#### 2.1.3. Total internal reflection fluorescence (TIRF) microscope

The objective-type TIRF microscope uses a 488 nm diode-laser with a maximum power of 80 mW (PhoxX 488-80, Omicron-Laserage, Germany). The laser is magnified by a telescope and cleaned up by a Fourier-filter with a diameter of 30 µm. To focus the laser into the objective’s BFP, a lens and a small, millimeter-sized mirror [31] are mounted on a closed-loop piezo-driven stage (M3-L, NewScale Technologies, USA) underneath the objective. The piezo-driven stage allows to precisely position the laser at a certain distance from the optical axis. Thus, the angle of incidence can be precisely tuned to achieve total internal reflection at the glass–water interface. Reflected excitation light is coupled out by a second small mirror that is also motorized by the same piezo stage as used for the other small mirror. Fluorescent light is collected by the objective lens and reflected by a dichroic mirror (F73-510, AHF Analysentechnik, Germany) toward the fluorescence detection path. A 160 mm tube-lens forms an image of the sample and an adjustable slit (Spalte SP 40, Owis, Germany) confines the field of view. The following unit is a color splitter, independently developed and similar in design to the one by Jiang *et al.* [32]. The fluorescent light is split into a “green” and “red” path by the usage of a dichroic mirror (F38-560, AHF Analysentechnik, Germany) and focuses both colors independently onto an electron multiplying CCD camera (iXon3, Andor Technology, UK).

### 2.2. Sample preparation

#### 2.2.1. Polymerization of microtubules

Microtubules were polymerized from a 30 µM tubulin solution that contained, 4 mM MgCl_2_, 1 mM GTP, and 5 % DMSO in BRB80 (80 mM PIPES/KOH pH 6.9, 1 mM MgCl_2_, 1 mM EGTA). For the fluorescence experiments, 10 % rhodamine-labeled tubulin was used. The solution was incubated for 30 min at 37 ^°^C. The polymerized microtubules were then centrifuged and suspended in a 0.1 % taxol–BRB80 solution.

#### 2.2.2. Microsphere functionalization

Polystyrene (PS) microspheres with a diameter of 590 nm (Bangs Laboratories, USA) were functionalized as described in Bugiel *et al.* [33] except that the GFP antibody was replaced with a nanobody: a GFP-binding-protein (GBP), which is a 13 kDa GFP-binding fragment derived from a llama single-chain antibody [34]. A protein solution of a truncated rat kinesin-1 motor protein (his6-rkin430-eGFP) was then mixed with the microsphere solution. The protein solution contained BRB80, 112.5 mM casein, 1 mM AMPPNP (a non-hydrolyzable analog of ATP ensuring a rigor kinesin-microtubule attachment), and an oxygen scavenger system (20 mM D-glucose, 20 µg/ml glucose oxidase, 8 µg/ml, and 0.1 % β-mercaptoethanol).

#### 2.2.3. Sample chamber preparation

The sample flow-cells were constructed of Parafilm ^®^ sandwiched between two cover slips (22 mm × 22 mm, # 1.5, Corning, and 18 mm × 18 mm, # 0, Menzel Gläser), which were cleaned and treated with chlorotrimethylsilane vapor under vacuum to generate a hydrophobic surface. The flow chamber was first filled with a solution containing anti-β-tubulin I antibodies (Sigma-Aldrich). For the microtubule bending experiment, we decreased the concentration of anti-β-tubulin about 1000×. Non-specific binding was prevented by incubating the flow chamber for 10 min with a 1 % aqueous solution of Pluronic F-127 (Sigma-Aldrich). Afterwards, the microtubule solution was flowed in. To bend a microtubule, the solution of functionalized microspheres was added after the microtubules were bound to the surface.

### 2.3. Image processing

#### 2.3.1. Background measurement

Inhomogeneous illumination of the sample is a common problem in microscopy and is accounted for by subtraction of a so-called background image. In IRM, the background strongly depends on the vertical position of the reference plane, i.e. the glass–water interface relative to the focal or imaging plane. Therefore, it is necessary to move the reference into the focal plane to measure a background image. To avoid any (static) sample of being considered as background, we moved the sample laterally, sufficiently fast, while recording a set of ≈400 images at a frame rate of 25 Hz. A median of this set is calculated and considered as background. This background image is subtracted from the IRM images (Fig. 3A). After subtraction, the IRM images show the interference contrast as a positive or negative signal. Note that the nominal values depend on the bit-depth of the recorded images and should be normalized by this depth if raw signal values need to be compared.

**Fig. 3:**
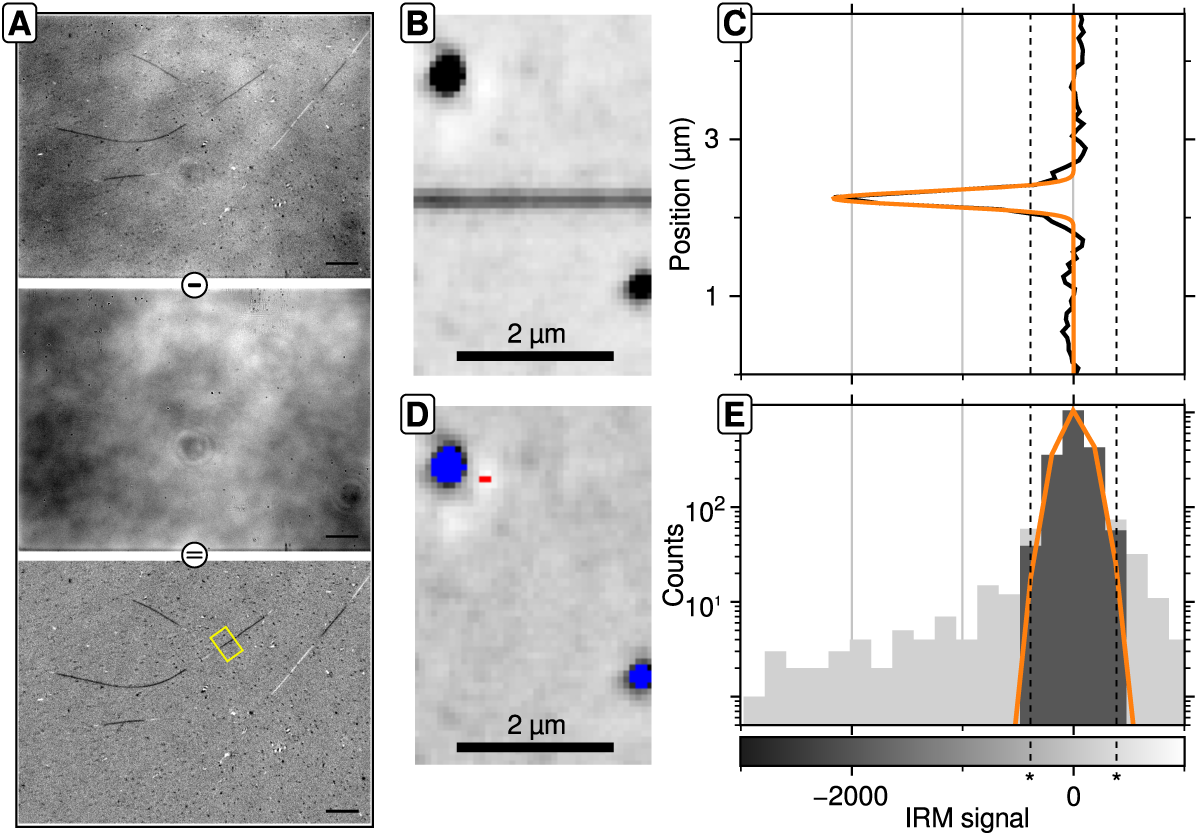
**Background subtraction and SNR measurement of a microtubule. (A)** Raw (top) and background (middle) IRM image of microtubules on a glass surface. Background subtraction leads to the image of the IRM signal (bottom). **(B)** Region of interest of the microtubule indicated in A. The black spots are impurities. **(C)** Median-intensity profile calculated along the microtubule-axis of B (black) and a Gaussian-fit (orange). **(D)** Residual noise image after subtraction of the Gaussian-fit shown in C from each column of the microtubule-image in B. Values above or below the threshold (vertical, dashed lines in C) are indicated in red or blue, respectively. **(E)** Histogram of the gray values in D (light gray bars). The image noise was calculated from the gray values within the threshold (dark gray bars) excluding signals from impurities. The threshold (*) was calculated as ±1.5× the interquartile range around the median gray level.

#### 2.3.2. Measuring the signal-to-noise ratio

To compare different IRM images it is necessary to use a standard signal-generating sample. Here, we used the contrast generated by a single microtubule. One way of measuring a microtubule’s signal is based on an intensity profile of a line across the microtubule. The difference between the highest and lowest intensity would be the signal. However, this approach is error-prone because the IRM signal depends on the probe’s height above the surface and potential irregularities as well as dirt on the glass surface. The latter, in particular, could result in large errors in the signal value.

Here, the signal of a microtubule was extracted from a Gaussian-fit to a median intensity profile calculated along the microtubule-axis. First, a part of a microtubule was cut out, using *Fiji’s* segmented-line in combination with the “Straighten” tool [35] (Fig. 3B). Then, the median intensity profile along the microtubule-axis was calculated (Fig. 3C). The microtubule signal, *I*_0_, was then extracted from a Gaussian-fit (*I* (*x*) = *I*_0_ exp (–(*x − x*_0_) ^2^ /2σ^2^) + *I*_offset_) to the median profile. The noise is calculated from a filtered residual image. An image of the IRM signal contains noise, whereby the major intrinsic contribution of noise is shot noise. Other contrast generating content, such as dirt, drift, or fluctuations in illumination intensity add to the total noise. To mainly extract the shot noise, a residual image was calculated by subtracting the Gaussian-fit from every column of a region of interest centered around a microtubule (Fig. 3D). A direct calculation of the standard deviation of the intensity distribution of the residual image usually overestimates the perceptual noise because dirt or irregularities in the glass surface increase the variance. To minimize the overestimation, we applied a threshold to the noise histogram. The threshold was set at *Q*_1_ − 1.5 IQR and *Q*_3_ + 1.5 IQR, where *Q*_1_ and *Q*_3_ are the first and the third quartile of the intensity distribution and IQR is its interquartile range (Fig. 3E). The resulting intensity distribution of the filtered residuals was then fitted to another Gaussian. Its standard deviation was taken as a measure for the noise. The ratio between the signal and the noise was then taken as the measure for the IRM signal quality or SNR.

## 3. Results & discussion

### 3.1. Optimizing the signal-to-noise ratio

The SNR of a microtubule increases with the amount of light reaching the camera. To investigate how the signal of a microtubule can be improved, we kept the exposure time constant at 0.04 s and consecutively increased the illumination light power by increasing the LED current until we saturated the CCD camera (Fig. 4A). The SNR increased in a power-law manner with an exponent 0.44 ± 0.03. This increase approximately follows the theoretical behavior in the shot noise limit with an exponent of 0.5. Deviations may be due to small non-linearities in the LED power-current relationship. In these experiments, the SNR did not saturate. Therefore, the pixel well-depth of our camera was limiting the SNR because higher LED currents saturated the camera. In the shot-noise limit, the SNR improves by averaging consecutive frames.

**Fig. 4:**
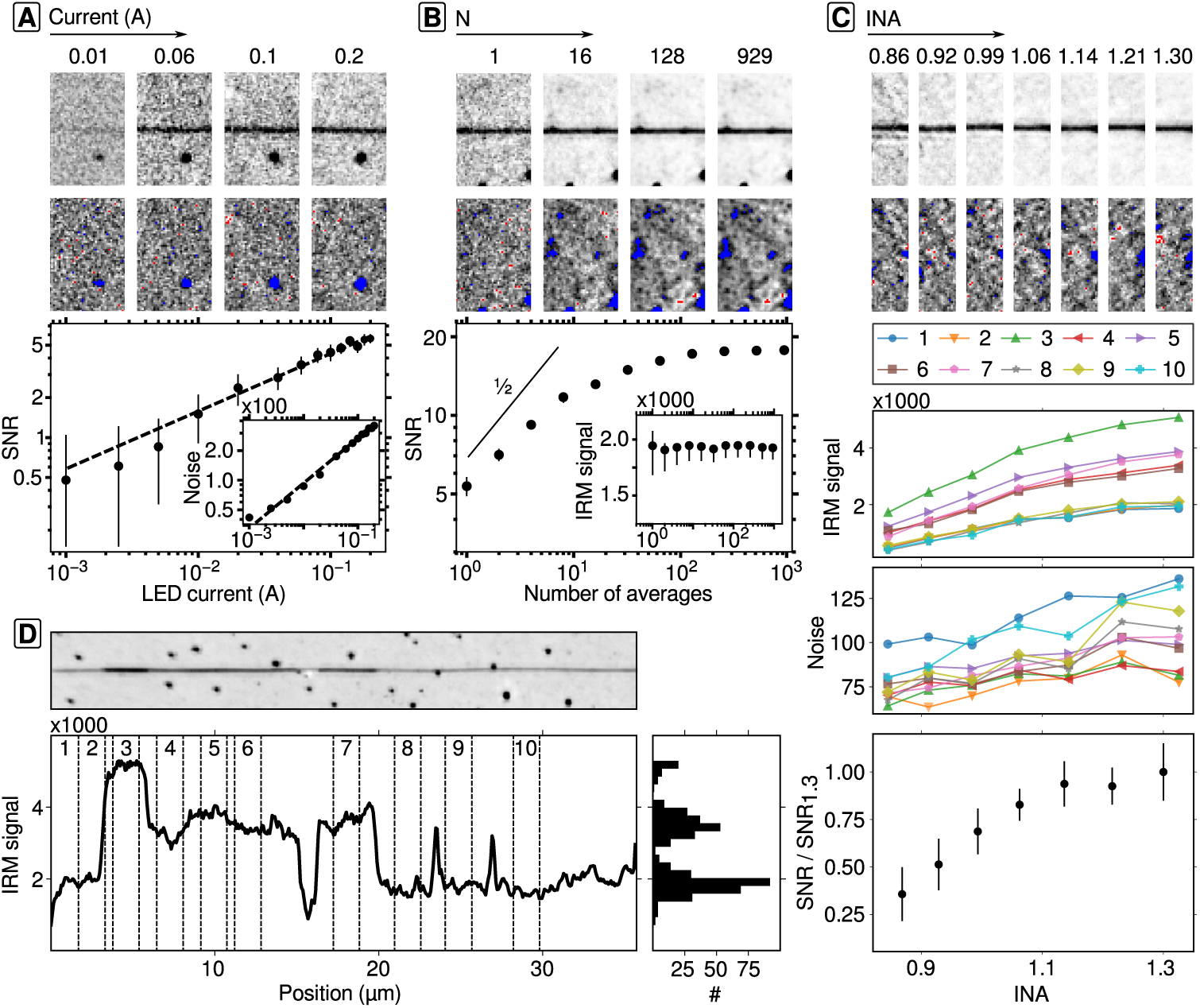
**Optimization of the IRM signal-to-noise ratio. (A)** Analyzed microtubule images (top) and the image noise (middle) at varying LED currents. The measured SNR is plotted against the applied LED current (bottom). A power-law fit (dashed line) showed near shot-noise limited imaging. Inset: Measured noise vs. LED current. The noise scaled with an exponent 0.46 ± 0.1. **(B)** Analyzed microtubule images (top) and the image noise (middle) for increasing number *N* of averaged frames. The SNR increases with the number of averaged frames, but reaches a plateau at large *N* (bottom) as irregularities dominate the noise. The solid line indicates a power-law slope with exponent 0.5. Inset: IRM signal vs. number of averages. The IRM signal of the microtubule was constant with respect to the number of averaged frames. **(C)** Top 2 panels: section 10 images of the microtubule shown in D with increasing INA and the respective image noise. Bottom panels: Legend of sections. Signal of all microtubule-sections (indicated in D) and the corresponding noise is plotted vs. INA. The averaged SNR is normalized by the SNR at INA = 1.3 at the very bottom. **(D)** An averaged image of a microtubule (*N* = 50), its line intensity profile (bottom left) and the corresponding histogram (bottom right). The analyzed microtubule-sections of C are indicated (dashed lines).

The SNR of a microtubule increased with the number of averaged frames, but was limited by other contrast-generating irregularities in the sample (Fig. 4B). We recorded a set of ≈900 images of a single microtubule at a frame rate of 25 Hz using an LED current of 0.2 A and measured its SNR as a funtion of the number of averaged frames. In one frame, a single microtubule had a SNR of ≈5.6. Upon averaging, the SNR doubled at *N* = 8 frames (SNR ≈ 11.4), but levelled off to ≈17.2 for *N* ≥ 128 frames. The saturation was due to the effective image noise becoming constant because the measured IRM signal of the microtubule was also constant (Fig. 4B, bottom, inset). While for a single frame, shot-noise was the dominant noise, the signal of contrast-generating irregularities surpassed the shot-noise for a high number of averages. The increase of the SNR is, therefore, limited by the cleanliness of the sample and the roughness of the glass cover slips. In analogy to the procedure reported for iSCAT imaging [24, 25], the background could be reduced by dynamic background image acquisition and subtraction.

The SNR could be further improved by using a high illumination numerical aperture (INA). By opening and closing the aperture iris, we varied the INA and, thereby, the LED image size in the BFP of the objective. To account for the change in average intensity on the CCD, we adjusted the LED current to a constant mean CCD intensity. We imaged one long microtubule, averaged 50 consecutive frames, and measured the signal and the image noise of 10 different microtubule sections (Fig. 4C, D). The signal of a microtubule-section and its image noise increased with increasing INA (Fig. 4C, middle). The resulting SNR of all microtubule-sections increased up to an INA of 1.14 and levelled off for higher INAs (Fig. 4C, bottom). Increasing the illumination angle improved the IRM signal, but also increased the image noise. The increase of the image noise might again be caused by an increasing visibility of irregularities on the glass surface. However, the increase of the IRM signal was not expected based on theory (compare Eq. (1) below). A quantitative comparison of the INA dependence for samples located directly at the glass surface (*h* = 0) or at varying heights did not account well for the increase in the SNR. The measurements might be explained by other effects, e.g. by empty-aperture, for which high-order spherical aberrations lead to an effective loss of NA [36]. A non-uniform scattering field of small molecules like microtubules, which was recently utilized to improve iScat contrast [37, 38], may also contribute to the effect even though the imaging NA was unchanged during our experiments.

IRM resolved bundled microtubules. Some microtubules (e.g. Fig. 4D) showed a patchy IRM signal. In IRM, differences in the signal could occur due to height differences of the specimen or different mass. In the former, changes in height typically occur gradually. In the latter, the signal changed in a stepwise fashion within the lateral resolution limit. The histogram of a line profile along the microtubule in Fig. 4D showed a discrete distribution. The discrete peaks suggests that several microtubules were bundled consistent with previous observations [39].

### 3.2. Quantitative 3D-IRM

To test whether the IRM signal can be used for precise 3D measurements of samples and calibration of the evanescent TIRF field, we used optical tweezers to bend a microtubule upward in a controlled and calibrated fashion (Fig. 5A). To this end, we trapped a microsphere that was functionalized to bind microtubules (see Methods 2.2.2), attached the microsphere to a microtubule end, and pulled the microtubule end upward by displacing the optical trap in 200-nm vertical steps using the deformable mirror (Fig. 1). Since the other microtubule end was bound to the glass surface, the microtubule bent with increasing applied force. The increasing vertical position of the microtubule caused a phase change between the reference and scattered field and, therefore, an amplitude change of the IRM signal. To quantify this amplitude change and microtubule deformation, we recorded the IRM and TIRF signals (Fig. 5B). Subsequently, we measured the line-intensity profiles along the microtubule and determined its 3D profile, *h*(*x*) (Fig. 5C, E). The microtubule height, *h*(*x*), as a function of lateral position, *x*, is related to the IRM intensity *I*(*x*) according to [20, 40]

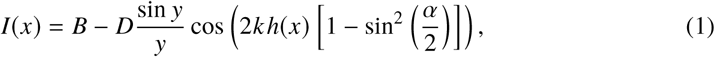

where *B* is the background gray-level intensity, which—after background subtraction—should be close to zero, *D* is the peak-to-peak interference amplitude, *k* = 2π*n*_w_ /λ is the wave number with the refractive index of water, *n*_w_, and the illumination wavelength, λ. The parmeter *y* = 2*kh*(*x*) sin^2^ α 2 models the objective’s point-spread-function, where α = arcsin *INA n*_w_ is the half angle of the INA cone. The interference amplitude, *D*, can also be related to the maximum iScat signal by *D* = 2*sr I*_inc_, with *I*_inc_ being the incident intensity, *r* is the reflectivity given by the refractive index mismatch between the glass–water interface and *s* is the scattering amplitude [22]. The microtubule itself can be modeled as a cantilevered beam of length *L* = *x*_*t*_ − *x*_0_, where *x*_*t*_ = 0 nm is the position of the trap and *x*_0_ is the position, at which the microtubule is attached to the surface. The beam equation in the small angle approximation is given by [41]

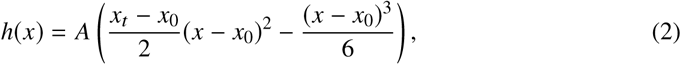

where *A* = *EI F* is the shape defining parameter given by the ratio of the flexural rigidity, i.e. the product of the elastic modulus *E* and the geometrical moment of inertia *I*, and the applied force *F*, which is provided by the optical trap.

**Fig. 5:**
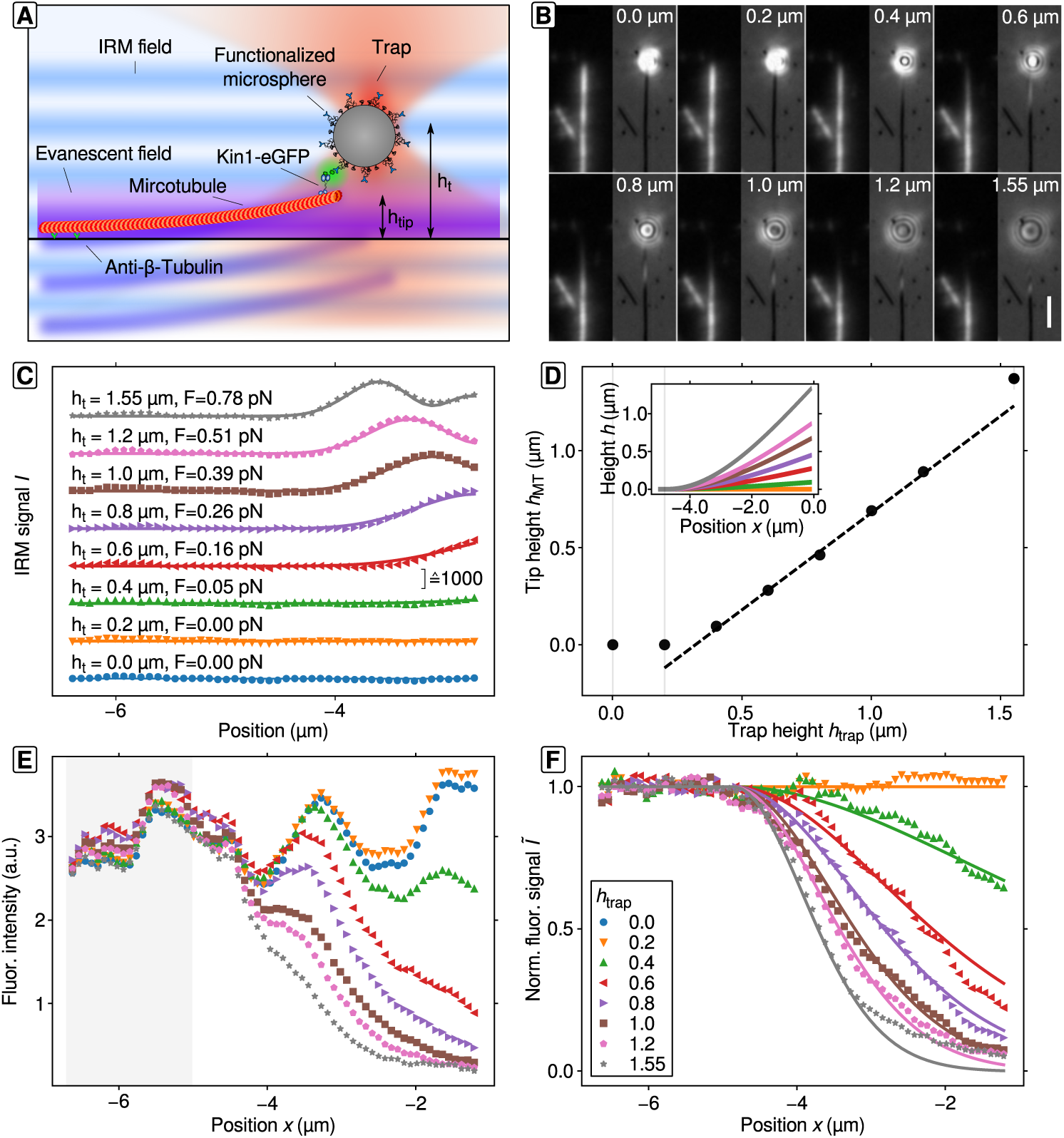
**IRM and evanescent field calibration using bent microtubules. (A)** Schematic drawing not to scale. A microsphere functionalized with non-motile kinesin-1–GFP motor proteins is trapped and attached to a loose microtubule end, which is pulled upward by the trap. **(B)** TIRF (red channel, left) and IRM images (right) of a bent microtubule at increasing trap heights. Scale bar: 2 µm. **(C)** IRM line profiles along the microtubule shown in B (symbols) and the global fit of Eq. (1) (solid lines). The trap height and axial force are indicated. For clarity, data points are offset vertically. **(D)** Weighted linear fit (dashed line) to the microtubule tip height *h*_MT_ (circles) plotted versus the trap height *h*_trap_ (error bars are SEM, gray lines). Inset: Calculated microtubule profile vs. height. **(E)** Fluorescence intensity line profiles along the microtubule shown in B for different tip heights. **(F)** Normalized fluorescence intensity line profiles (symbols). Profiles were normalized to the average intensity of the first 20 points and to the line profile at *h*_t_ = 0.0 µm (shaded area and blue circles in E, respectively). A global fit (solid lines) of the data to Eq. (3) determined the depth of the evanescent field.

A global non-linear least-squares fit using Eqs. (1) and (2) to the microtubule IRM profiles (Fig. 5C) resulted in the height of the microtubule with respect to its lateral position along the microtubule axis. The best fit parameters are listed in Table 1. Because axial optical tweezers force measurements require significant background correction, which make small-force measurements unreliable in particular for large trap-surface variations [14, 42], we estimated the force *F* that was acting on the microtubule based on its flexural rigidity of 20 × 10^−24^ Nm^2^ [41, 43]. Equation (2) allowed us to determine the height of the microtubule tip *h*_MT_ = *h*(*x*_*t*_) and the corresponding height-profile of the microtubule (Fig. 5D). A linear fit of the microtubule tip height *h*_MT_ with respect to the height of the trap *h*_trap_ showed the expected slope of unity and an offset of (–320 ± 10) nm. The offset roughly corresponds to the microsphere radius (≈295 nm) plus the linker size (≈35 nm). The axial displacement of the microsphere within the trap in the nanometer range was negligible compared to the tip height in the micrometer range (with the maximum force of *F*_*z*_ ≈0.8 pN and a trap stiffness κ_z_ in the vertical direction of (0.05 ± 0.01) pN/nm, the displacement was Δ*z* = *F*_*z*_ /κ_*z*_ ≈ (16± 4) nm). The agreement between the IRM-based microtubule tip height and predefined optical trap height confirms that the IRM signal can be used for quantitative 3D measurements.

**Table 1:** Fit parameters of the global fit to the IRM and TIRF intensity profiles. The IRM global fit had *B* = 0, *x*_0_, *D*, and INA as common parameters for all IRM line profiles and a set of parmeters, *A*, that corresponded to the force values *F* = *EI/A* for each individual line profile. The global fit to all normalized TIRF line profiles had the evanescent field depth δ as common parameter.

| **IRM fit** |  |  |
| --- | --- | --- |
| $x_0$ ($\mu\text{m}$) | $-4.72 \pm 0.02$ | |
| $D$ | $979 \pm 5$ | |
| $INA$ | $1.15 \pm 0.01$ | |
| $h_{\text{trap}}$ ($\mu\text{m}$) | $A$ ($\times 10^{-3} \mu\text{m}^{-2}$) | $F$ (pN) |
| 0.0 | $0.0 \pm 314$ | $0.00 \pm 6$ |
| 0.2 | $0.0 \pm 3347$ | $0.00 \pm 66$ |
| 0.4 | $2.7 \pm 0.4$ | $0.05 \pm 0.01$ |
| 0.6 | $8.0 \pm 0.3$ | $0.16 \pm 0.01$ |
| 0.8 | $13.2 \pm 0.4$ | $0.26 \pm 0.01$ |
| 1.0 | $19.7 \pm 0.6$ | $0.39 \pm 0.01$ |
| 1.2 | $25.4 \pm 0.7$ | $0.51 \pm 0.01$ |
| 1.55 | $39.0 \pm 1.2$ | $0.78 \pm 0.02$ |
| **TIRF fit** |  |  |
| $\delta$ (nm) | $165 \pm 4$ | |

As a further quantitative control for the accuracy of the IRM-based height profiles, we tested whether the height profiles are consistent with the exponential decay of the TIRF field. In addition, we can use the profiles to measure the evanescent field depth. The TIRF intensity of the profiles (Fig. 5B, E) depend on both the fluorophore labelling density and the fluorophore height above the surface. To account for the inhomogeneous labeling and photo-bleaching, we normalized the TIRF intensity profiles (Fig. 5F). The normalized profiles were consistent with an exponentially decreasing TIRF intensity Ĩ with increasing distance from the glass–water interface

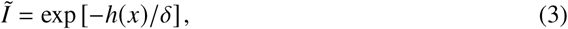

where *δ* is the evanescent field depth. With the known shape of the microtubule, the decay of fluorescence intensity solely depends on the depth of the evanescent field, *δ*. A global fit of the normalized fluorescence intensity profiles to Eq. (3) with *h* = *h* (*x, x*_0_, *A*) and the evanescent field depth as common parameter resulted in *δ* = (165 ± 4) nm, which is in good agreement with our experimental design. In summary, the microtubule IRM signal quantitatively agreed in 3D with both optical trap and TIRF based height determination.

## 4. Conclusion

Using LED-IRM, we were able to measure the 3D profile of a single microtubule with high contrast and precision. To characterize the quality of the signal, we developed a robust method to measure the SNR of a single microtubule. Our method accounts for overestimation of the microtubule signal due to contrast generating structures that might lie underneath the microtubule. It also accounts for overestimation of image noise by avoiding high-contrast signals in the residual image that result from other contrast generating structures. LED-based IRM is limited by shot noise as the SNR scales with the available light and is, thus, limited by the electron well-depth of the camera. The SNR of a single microtubule in one frame was about 5.6. This SNR is comparable to the one achieved with DIC (SNR ≈ 3.4 [2]). The value is also consistent with recent work using IRM and comparing it to different microscopy techniques [28]. IRM interference fringes give information about the height of a specimen. Although the quantitative interpretation of the fringes, in general, is a non-trivial task, especially for arbitrary objects [40], in cases where the structure of interest is simple, such as a bent microtubule, it is possible to determine the axial position of an object and thus, the 3D profile with high precision on the order of a few tens of nanometers. This knowledge we used to determine the depth of the evanescent field of the TIRF microscope.

LED-based IRM is a cheap and simple, high-contrast 3D microscopy method, which can be integrated easily into existing optical setups providing a viable alternative to commonly used microscopy techniques. IRM does not require expensive nor complicated optical elements like lasers or polarizing optics. Especially for single-molecule force spectroscopy using optical tweezers, IRM has considerable advantages compared to DIC microscopy. First, there is no trapping power loss (DIC prisms may reduce the trapping power by up to 10%). Second, whereas the SNR of a microtubule is comparable to the one achieved in DIC [2], IRM outcompetes DIC with its simple optical design, cheap components and its orientation-independent contrast. Third, in contrast to DIC, IRM does not restrict the optical tweezers design to a linearly polarized trapping laser. Therefore, any polarization state like circularly polarized light is possible allowing the use of an optical microprotractor, torsion balance, or torque wrench [17, 44]. Also, because illumination and detection is done via the objective lens, the sample is freely accessible from the top for other manipulation or imaging techniques. Furthermore, LED-based illumination provides long-term intensity stability, if a sufficiently stable power supply and heat sink are used. In contrast to laser-based illumination, where long coherence lengths can cause etalon fringes, i.e. “unwanted” interference effects, light emitted from an LED has a coherence length

## Funding

Deutsche Forschungsgemeinschaft (DFG; Emmy Noether Program and CRC 1101, Project A04); European Research Council (ERC Starting Grant 2010, Nanomech 260875); Technische Universität Dresden; and Universität Tübingen.

## Contributions

S.S. designed, constructed, and assembled the instrument, prepared samples for the SNR measurements and performed all experiments. M.K.A. prepared the quantitative IRM experiments, i.e. functionalization of microspheres and flow cell preparation. G.H. helped in the design and assembly of the TIRF microscope. S.S. and E.S. conceived and designed the experiments, analyzed the data, and wrote the manuscript.

## Acknowledgements

The authors thank U. Rothbauer for providing the GFP-binding protein and M. Bugiel, A. Jannasch, M. Mahamdeh and Q. T. Davis for discussions and critical reading of the manuscript. This work was done by use of free and open software. S. S. wants to acknowledge the following projects in particular: Python [45], Scipy [46], matplotlib [47] and lmfit [48].

